# μDamID: a microfluidic approach for imaging and sequencing protein-DNA interactions in single cells

**DOI:** 10.1101/706903

**Authors:** Nicolas Altemose, Annie Maslan, Andre Lai, Jonathan A. White, Aaron M. Streets

## Abstract

Genome regulation depends on carefully programmed protein-DNA interactions that maintain or alter gene expression states, often by influencing chromatin organization. Most studies of these interactions to date have relied on bulk methods, which in many systems cannot capture the dynamic single-cell nature of these interactions as they modulate cell states. One method allowing for sensitive single-cell mapping of protein-DNA interactions is DNA adenine methyltransferase identification (DamID), which records a protein’s DNA-binding history by methylating adenine bases in its vicinity, then selectively amplifies and sequences these methylated regions. These interaction sites can also be visualized using fluorescent proteins that bind to methyladenines. Here we combine these imaging and sequencing technologies in an integrated microfluidic platform (μDamID) that enables single-cell isolation, imaging, and sorting, followed by DamID. We apply this system to generate paired single-cell imaging and sequencing data from a human cell line, in which we map and validate interactions between DNA and nuclear lamina proteins, providing a measure of 3D chromatin organization and broad gene regulation patterns. μDamID provides the unique ability to compare paired imaging and sequencing data for each cell and between cells, enabling the joint analysis of the nuclear localization, sequence identity, and variability of protein-DNA interactions.

## Introduction

Complex life depends on the protein-DNA interactions that constitute and maintain the epigenome, including interactions with histone proteins, transcription factors, DNA (de)methylases, and chromatin remodeling complexes, among others. These interactions enable the static DNA sequence inside the nucleus to dynamically execute different gene expression programs that shape the cell’s identity and behavior. Methods for measuring protein-DNA interactions have proven indispensable for understanding the epigenome, though to date most of this knowledge has derived from experiments in bulk cell populations. By requiring large numbers of cells, these bulk methods can fail to capture critical epigenomic processes that occur in small numbers of dividing cells, including processes that influence embryo development, developmental diseases, stem cell differentiation, and certain cancers. By averaging together populations of cells, bulk methods also fail to capture important epigenomic dynamics occurring in asynchronous single cells during differentiation or the cell cycle. Because of this, bulk methods can overlook important biological heterogeneity within a tissue. It also remains difficult to pair bulk biochemical data with imaging data, which inherently provide information in single cells, and which can reveal the spatial location of protein-DNA interactions within the nuclei of living cells. These limitations underline the need for high-sensitivity single-cell methods for measuring protein-DNA interactions.

Most approaches for mapping protein-DNA interactions rely on immunoaffinity purification, in which protein-DNA complexes are physically isolated using a high-affinity antibody against the protein, then purified by washing and de-complexed so the interacting DNA can be amplified and measured. The most widely used among these methods is chromatin immunoprecipitation with sequencing (ChIP-seq; Barski et al. 2007, Johnson et al. 2007), which has formed the backbone of several large epigenome mapping projects (Celniker et al. 2009; ENCODE Consortium 2012; Kundaje et al. 2015). One drawback of ChIP-seq is that the protein-DNA complex, which is often fragile, must survive the shearing or digestion of the surrounding DNA, as well as several intermediate washing and purification steps, in order to be amplified and sequenced. This results in a loss of sensitivity, especially when using a small amount of starting material. More recent immunoaffinity-based methods have reduced the high input requirements of ChIP-seq, but they recover relatively few interactions in small numbers of cells or single cells (Wu et al. 2012, Shen et al. 2014, Jakobsen et al. 2015, Rotem et al. 2015, Zhang et al. 2016, Skene et al. 2018, Harada et al. 2018, Kaya-Okur et al. 2019, Carter et al. 2019, Grosselin et al. 2019).

An alternative method for probing protein-DNA interactions, called DNA adenine methyltransferase identification (DamID), relies not on physical separation of protein-DNA complexes (as in ChIP-seq), but on a sort of ‘chemical recording’ of protein-DNA interactions onto the DNA itself, which can later be selectively amplified (Figure 1a; van Steensel and Henikoff 2000, Vogel et al. 2007). This method utilizes a small enzyme from *E. coli* called DNA adenine methyltransferase (Dam). When genetically fused to the protein of interest, Dam deposits methyl groups near the protein-DNA contacts at the N6 positions of adenine bases (^m6^A) within GATC sequences (which occur once every 270 bp on average across the human genome). That is, wherever the protein contacts DNA throughout the genome, ^m6^A marks are left at GATC sites in its trail. These ^m6^A marks are highly stable in eukaryotic cells, which do not tend to methylate (or demethylate) adenines (O’Brown et al. 2019). Dam expression has been shown to have no discernable effect on gene expression in a human cell line, and its ^m6^A marks were shown to be passed to daughter cells, halving in quantity each generation after Dam is inactivated (Park et al. 2018). These properties allow even transient protein-DNA interactions to be recorded as permanent, biologically orthogonal chemical signals on the DNA.

**Figure 1.**
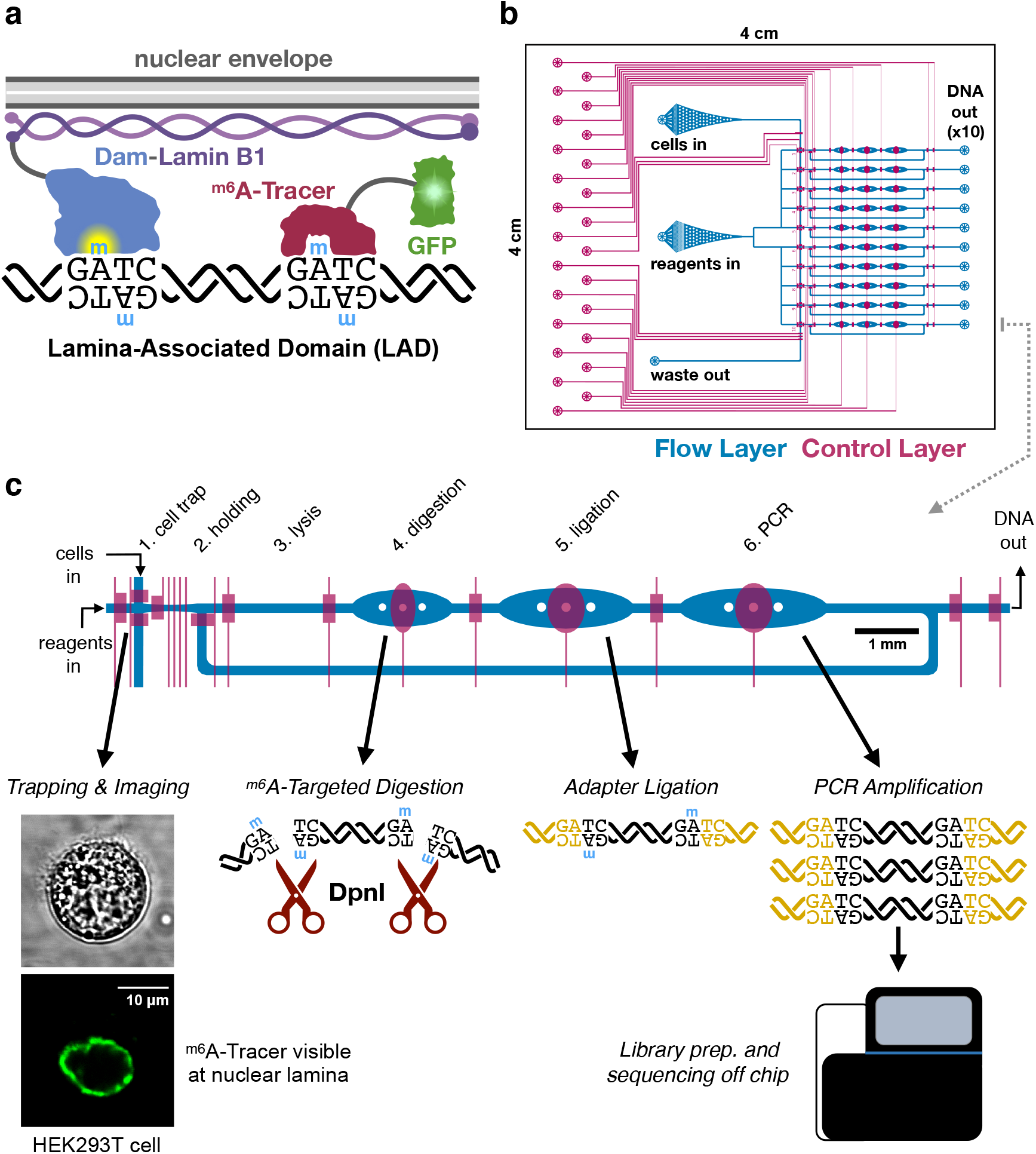
μDamID device design and function. **a)** overview of DamID (van Steensel and Henikoff 2000) and ^m6^A-Tracer (Kind et al. 2013) technologies applied to study interactions between DNA and nuclear lamina proteins. **b**) the overall design of the 10-cell device, showing the flow layer (blue, where cells and reagents enter channels) and the control layer (red, where elastomeric valves overlap the flow layer to control the flow of liquids). **c**) a closer view of one lane explaining the DamID protocol and the function of each chamber of the device. 10 cells are trapped, imaged, and selected serially, one per lane, then all 10 cells are lysed and processed in parallel.

DamID reads out these chemical recordings of protein-DNA interactions by specifically amplifying and then sequencing fragments of DNA containing the interaction site. First, genomic DNA is purified and digested with DpnI, a restriction enzyme that exclusively cleaves G ^m6^ATC sites (Figure 1). Then, universal adapters are ligated onto the fragment ends to allow for amplification using universal primers. Only regions with a high density of ^m6^A produce DNA fragments short enough to be amplified by Polymerase Chain Reaction (PCR) and quantified by microarray or high-throughput sequencing (Wu et al. 2016). DamID has been used to explore dynamic regulatory protein-DNA interactions such as transcription factor binding (Orian et al. 2003) and RNA polymerase binding (Southall et al. 2013) as well as protein-DNA interactions that maintain large-scale genome organization. One frequent application of DamID is to study large DNA domains associated with proteins at the nuclear lamina, near the inner membrane of the nuclear envelope (Pickersgill et al. 2006, Guelen et al. 2008, reviewed by van Steensel and Belmont 2017). Because DamID avoids the limitations of antibody binding, physical separations, or intermediate purification steps, it lends itself to single-cell applications. Recently, DamID has been successfully applied to sequence lamina-associated domains (LADs) in single cells in a one-pot reaction, recovering hundreds of thousands of unique fragments per cell (Kind et al. 2015).

While DamID maps the sequence positions of protein-DNA interactions throughout the genome, the spatial location of these interactions in the nucleus can play an important role in genome regulation (Bickmore and van Steensel 2013). A recent technique demonstrated the ability to specifically label and visualize protein-DNA interactions using fluorescence microscopy, revealing their spatial location within the nucleus in live cells (Kind et al. 2013). Visualization requires co-expression of a different fusion protein called ^m6^A-Tracer, which contains green fluorescent protein (GFP) and a domain that binds specifically to methylated GATC sites. This imaging technology has been applied to visualize the dynamics of LADs within single cells (Kind et al. 2013). Many recent efforts have aimed to measure chromatin organization in single cells, to better understand the heterogeneity of cells within tissues and the biological underpinnings of their gene expression states (reviewed by Kelsey et al. 2017). Both imaging and sequencing protein-DNA interactions can provide useful single-cell epigenomic information, but despite recent advances in single-cell sequencing technologies, it remains fundamentally difficult to track individual cells and pair their sequencing data with other measurements such as imaging. Pairing imaging and sequencing data could be applied to study, for example, how the dynamic remodeling of chromatin proteins across the genome in developing cells relates to the localization of those proteins in the nucleus.

Here we aimed to pair DamID with ^m6^A-Tracer imaging to produce coupled imaging and sequencing measurements of protein-DNA interactions in the same single cells. To achieve this, we engineered an integrated microfluidic device that enables single-cell isolation, imaging, selection, and DamID processing, which we call “μDamID.” We applied our device to image and map nuclear lamina interactions in a transiently transfected human cell line co-expressing ^m6^A-Tracer, and we validated our measurements against bulk DamID data from the same cell line as well as other human cell lines (Lenain et al. 2017, Kind et al. 2015). We discuss the advantages and potential applications of our device as well as future improvements to this system.

## Results and Discussion

### Design and operation of a microfluidic device with valve-actuated active cell traps

We designed and fabricated a polydimethylsiloxane (PDMS) microfluidic device with integrated elastomeric valves to facilitate the various reaction stages of the DamID protocol in a single liquid phase within the same device (Figure1). The device is compatible with high-magnification imaging on inverted microscopes, enabling imaging prior to cell lysis. Each device was designed to process 10 cells in parallel, each in an individual reaction lane fed from a common set of inlets. Valves are controlled by pneumatic actuators operated electronically via a programmable computer interface (White and Streets 2018).

Device operation was modified from our previous single-cell RNA sequencing platform (Streets et al. 2014). A suspension of single cells is loaded into the cell inlet (Figure 1b) and cells are directed towards a trapping region by a combination of pressure-driven flow and precise peristaltic pumping. As a cell crosses one of the 10 trapping regions, valves are actuated to immobilize the cell for imaging (Supplementary Figure 1). The cell is imaged by confocal fluorescence microscopy to visualize the localization of ^m6^A-Tracer, and after image acquisition, the user can choose whether to select the cell for DamID processing, or to reject it and send it out the waste outlet (Figure 1b).

Selected cells are injected from the trapping region into a holding chamber using pressure-driven flow from the reagent inlet (Figure 1b, Supplementary Figure 1). Once all 10 holding chambers are filled with imaged cells, processing proceeds in parallel for all 10 cells by successively adding the necessary reagents for each step of the single-cell DamID protocol (Kind et al. 2015) and dead-end filling each of the subsequent reaction chambers. Reaction temperatures are controlled by placing the device on a custom-built thermoelectric control unit for dynamic thermal cycling. Enzymes are heat inactivated between each step (Kind et al. 2015) and a low concentration of mild detergent was added to all reactions to prevent enzyme adhesion to PDMS (Streets et al. 2014).

Figure 1 shows a schematic of the microfluidic processing work flow. In the first reaction stage, a buffer containing detergent and proteinase pushes the cell into the lysis chamber, where its membranes are lysed and its proteins, including ^m6^A-Tracer, are digested away. Next, a DpnI reaction mix is added to digest the genomic DNA at Dam-methylated GATC sites in the digestion chamber. Then, a mix of DamID universal adapter oligonucleotides and DNA ligase is added to the ligation chamber. Finally, a PCR mix is added containing primers that anneal to the universal adapters is added and all valves within the lane are opened, creating a 120 nl cyclic reaction chamber. Contents are thoroughly mixed by peristaltic pumping around the reaction ring, then PCR is carried out on-chip by thermocycling. Amplified DNA is collected from each individual lane outlet, and sequencing library preparation is carried out off-chip.

### Application to map lamina-associated domains in a human cell line

We evaluated the performance of this platform by mapping the sequence and spatial location of lamina-associated domains in a human cell line, allowing us to compare our data to previously published LAD maps from DamID experiments in human cell lines (Kind et al. 2015, Lenain et al. 2017). LADs are large (median 500 kb) and comprise up to 30% of the genome in human cells (Guelen et al. 2008). LADs serve both a structural function, acting as a scaffold that underpins the three-dimensional architecture of the genome in the nucleus, and a regulatory function, as LADs tend to be gene-poor, more heterochromatic, and transcriptionally less active (reviewed by van Steensel and Belmont 2017 and Buchwalter et al. 2018). ^m6^A-Tracer has previously been applied to visualize LADs, which appear as a characteristic ring around the nuclear periphery in confocal fluorescence microscopy images (Kind et al. 2013; Figure 1c).

We carried out experiments in HEK293T cells for their ease of growth, transfection, suspension, and isolation. To enable rapid expression of Dam and ^m6^A-Tracer transgenes, we transiently transfected cells with DNA plasmids containing genes for a drug-inducible Dam-LMNB1 fusion protein as well as constitutively expressed ^m6^A-Tracer. We then induced Dam-LMNB1 expression, optimizing the expression times to maximize the proportion of cells with fluorescent laminar rings (Figure 1c). Because transient transfection yields a heterogeneous population of cells, each with potentially variable copies of the transgenes, it was important for us to be able to image cells and select only those with visible laminar rings, which were more likely to have the correct expression levels, and which were unlikely to be in the mitosis phase of the cell cycle. This kind of complex sorting would not be possible with sorting methods like fluorescence-activated cell sorting (FACS) but is straight-forward in our microfluidic platform.

In addition to processing Dam-LMNB1 cells, we transfected cells with the Dam gene alone, not fused to LMNB1, to provide a negative control demonstrating where the unfused Dam enzyme would mark the genome if not tethered to the nuclear lamina (Vogel et al. 2007). This control is useful for estimating the background propensity for each genomic region to be methylated, since Dam preferentially methylates more accessible regions of the genome, including gene-rich regions (Singh and Klar 1992, Lenain et al. 2017, Aughey et al. 2018). We selected Dam-only cells that had high fluorescence levels across the nucleus and did not appear mitotic. We also performed DamID in bulk transiently transfected HEK293T cells for validation (Vogel et al. 2007). We used a mutant of Dam (V133A; Elsawy and Chahar 2014), which is predicted to have weaker methylation activity than the wild-type allele on unmethylated DNA, to reduce background methylation. We performed bulk DamID experiments comparing the mutant and wild-type alleles and found that the V133A mutant allele provides more than twofold greater signal-to-background compared to the wild-type allele (Supplementary Figure 2). We also performed RNA sequencing in bulk cells that were untreated or transfected with Dam-only, Dam-LMNB1, or ^m6^A-Tracer, and we found only two differentially expressed genes (Supplementary Figure 3). This corroborates similar published findings by others showing that Dam expression and adenine methylation have little or no effect on gene expression in HEK293T cells (Park et al. 2018).

We ran three devices containing 25 imaged cells total, with empty lanes left as negative controls, which did not yield sequenceable quantities of DNA. From these, we selected 18 cells total for multiplexed sequencing, including 15 Dam-LMNB1 cells and 3 Dam-only cells, to achieve a desired level of coverage per cell. Selection was based on image quality and initial DNA quantification data from each sample (see Methods). We included one anomalous Dam-LMNB1 cell that appeared to have high fluorescence in the nuclear interior, predicting that it might have higher background DamID coverage in non-LAD regions (cell #7). After sequencing, we excluded 3 Dam-LMNB1 cells containing a high fraction of sequencing reads mapping to the transfected plasmids (Supplementary Figure 4); the 15 remaining cells had less than 5% of mapped reads mapping to plasmid DNA. For these 15 remaining cells, we obtained a median of roughly 600,000 raw reads per cell (range 300k– 2.7M), covering a median of 110,000 unique DpnI fragments per cell (37k – 370k), in line with previous DamID results from single cells (Kind et al. 2015).

### μDamID sequencing data recapitulate existing LAD maps

To assess whether our single-cell μDamID sequencing data provide accurate measurements of lamina-associated domains, we first compared our single-cell results to those we obtained from bulk DamID in the same cell line. DamID results are reported as a difference or log ratio between the observed coverage from Dam-LMNB1 expressing cells and the expected coverage from background, estimated using coverage from Dam-only expressing cells (see Methods). This measure is reported within fixed 250 kb bins across the genome, which is half the median length of known LADs in the genome (Kind et al. 2015). By aggregating the data from 11 Dam-LMNB1 expressing cells passing filters and excluding the anomalous cell #7, we found excellent correspondence with the bulk data obtained from millions of cells (Figure 2a), with a Pearson correlation of 0.85 across all bins in the genome. To ensure normalization is not inflating the correlation, we compared aggregate single-cell raw read coverage to bulk raw read coverage and observed a genome-wide correlation of 0.89 (Figure 2b).

**Figure 2.**
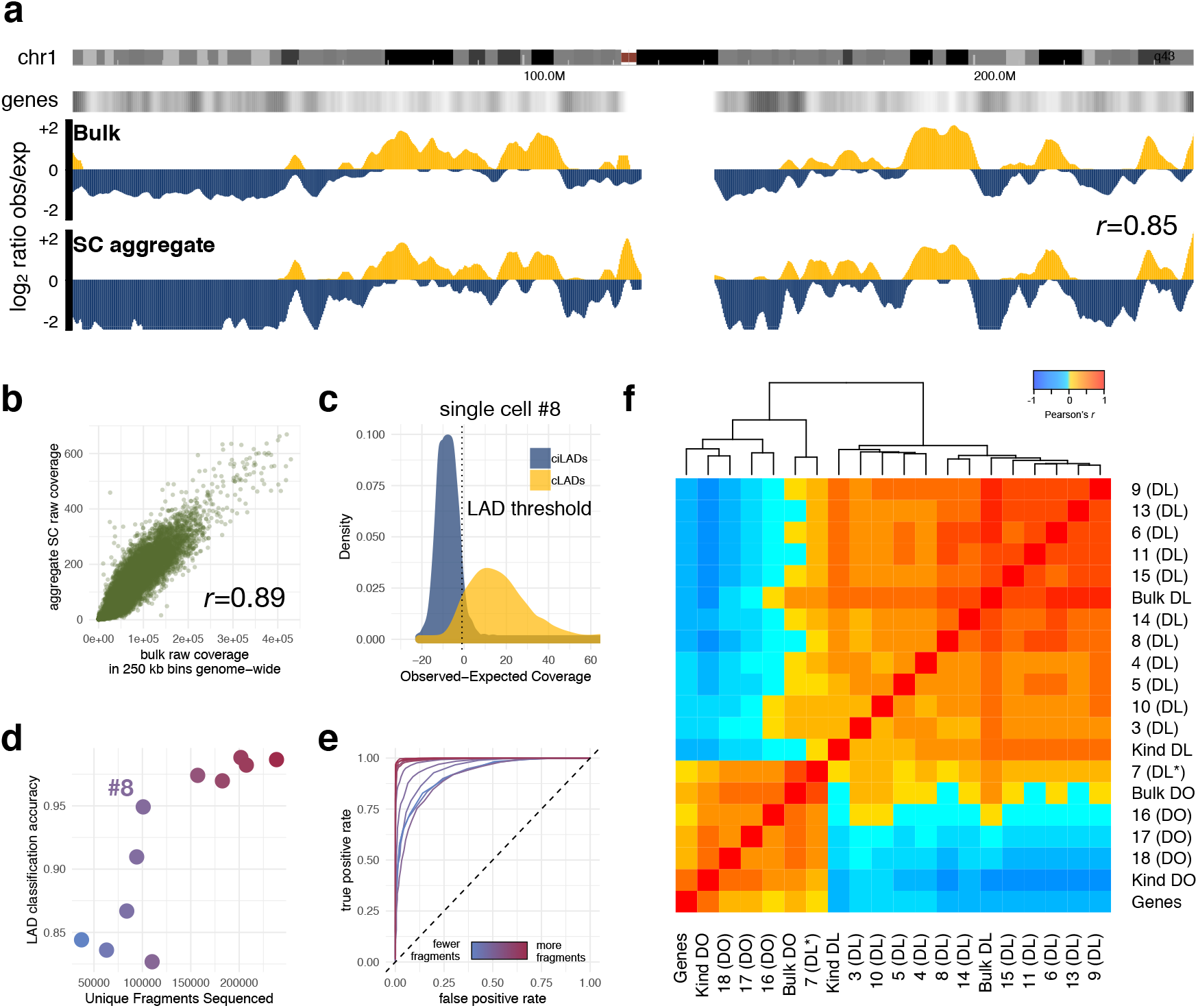
validation of μDamID sequencing data. (**a**) comparison of bulk DamID sequencing data and aggregate single-cell sequencing data across all of human chromosome 1. log_2_ ratios represent the ratio of Dam-LMNB1 sequencing coverage to normalized bulk Dam-only sequencing coverage. Positive values (gold) represent regions associated with the nuclear lamina, which tend to have lower gene density (second track from top). The Pearson correlation between bulk and aggregate single-cell data across all 250-kb bins in the genome is 0.85. (**b**) scatterplot comparing raw sequencing coverage in bulk and single cell samples (aggregated). (**c**) normalized coverage distribution in one single cell expressing Dam-LMNB1 (cell #8) in positive and negative control sets (cLADs, gold, and ciLADs, blue). The threshold that distinguishes these sets with maximal accuracy is shown as a dotted line. (**d**) The maximum control set classification accuracy for each of 11 Dam-LMNB1 cells versus the number of unique DpnI fragments sequenced for each cell (also indicated by colors). Cell #8, the sample with median accuracy plotted in **c**, is labeled. (**e**) Receiver-Operator Characteristic curves for all 11 cells, colored by the number of unique DpnI fragments sequenced. (**f**) pairwise Pearson correlation heatmap for raw sequencing coverage in 250 kb bins genome-wide, with dendrogram indicating hierarchical clustering results. Numbers indicate cell numbers. DL = Dam-LMNB1. DO = Dam-only. Genes = number of Refseq genes in each bin. Kind = aggregated single-cell data from Kind et al. 2015. Bulk = bulk HEK293T DamID data from this study. *anomalous Dam-LMNB1 cell (#7) with high ^m6^A-tracer signal in the nuclear interior.

We next computed pairwise correlations between the raw coverage for all single cells with each other, with the bulk data, with aggregated published single-cell DamID data (from Kind et al. 2015), and with the number of annotated genes in each 250 kb bin genome-wide. We performed unsupervised hierarchical clustering on these datasets and produced a heatmap of their pairwise correlations (Figure 2f). We found that the 3 Dam-only single cells cluster with each other, with the bulk Dam-only data, with the Kind et al. Dam-only data, and with the number of genes, as expected. The 11 Dam-LMNB1 cells cluster separately with each other, with the bulk Dam-LMNB1 data, and with the Kind et al. Dam-LMNB1 data. The anomalous cell #7 shows correlations with both the Dam-only and Dam-LMNB1 clusters, appearing intermediate between them (Figure 2f). This illustrates that our single-cell Dam-LMNB1 and Dam-only cells can be distinguished given their sequencing data alone, and they associate as expected with published data, with our bulk data, and with annotated gene density, further confirming that these sequencing data are measuring meaningful biological patterns in single cells. The anomalous cell #7 can also be distinguished by sequencing data alone, since its data correlate with both the Dam-only and Dam-LMNB1 cell data.

### μDamID enables accurate LAD calling within single cells

In order to define LADs across the genome within single cells, we trained a simple classifier on a set of stringent positive and negative controls: regions confidently known to be lamina-associated or not lamina associated based on bulk DamID data from our study and others (Lenain et al. 2017; see Methods). Positive controls consist of 250 kb bins across the genome that were previously annotated in other human cell lines and confirmed with bulk DamID in our own cell line to be consistently associated with the nuclear lamina (referred to as constitutive LADs, or cLADS). Negative controls were similarly determined using prior bulk data to be consistently not associated with the nuclear lamina (referred to as constitutive inter-LADs, or ciLADS). These stringent control sets constitute roughly 10% of the genome each.

For each single Dam-LMNB1 cell, we computed the distribution of its normalized sequencing coverage in bins from the positive and negative control regions (Figure 2c), with the expectation that ciLADs have little or no coverage and the cLADs have high coverage. Given these control distributions, we chose a coverage threshold to maximally separate the known cLADs and ciLADs. Across the 11 Dam-LMNB1 cells, we determined thresholds that distinguish the known cLADs and ciLADs with a median accuracy of 96% (range 83-99%), which correlates positively with the number of unique DpnI fragments sequenced per cell (Figure 2d). We also plotted receiver operating characteristic (ROC) curves for each cell, showing the empirical tradeoff between false positive and false negative LAD calls at varying thresholds (Figure 2e).

After choosing a threshold for each cell to maximize classification accuracy between the control sets, we applied these thresholds to make binary LAD classifications across the rest of the genome. At each bin in the genome, we counted the number of Dam-LMNB1 cells in which that bin was classified as an LAD (out of 11 total cells). As expected, bins belonging to the cLAD control sets are classified as LADs in almost all 11 of the cells while bins belonging to the ciLAD control sets are classified as LADs in almost none of the cells (Figure 3a-b). The intermediate bins (called as LADs in 4 to 7 cells), appearing to be lamina associated in only a subset of cells, are likely to contain regions that are variably associated with the lamina, differing from cell to cell, or possibly even dynamically moving between the lamina and the nuclear interior within the same cell over time (Kind et al. 2015). Single-cell data provide a unique opportunity to observe and measure this variability in chromatin organization between cells, enabling the identification of these variable LADs within a population of cells.

**Figure 3.**
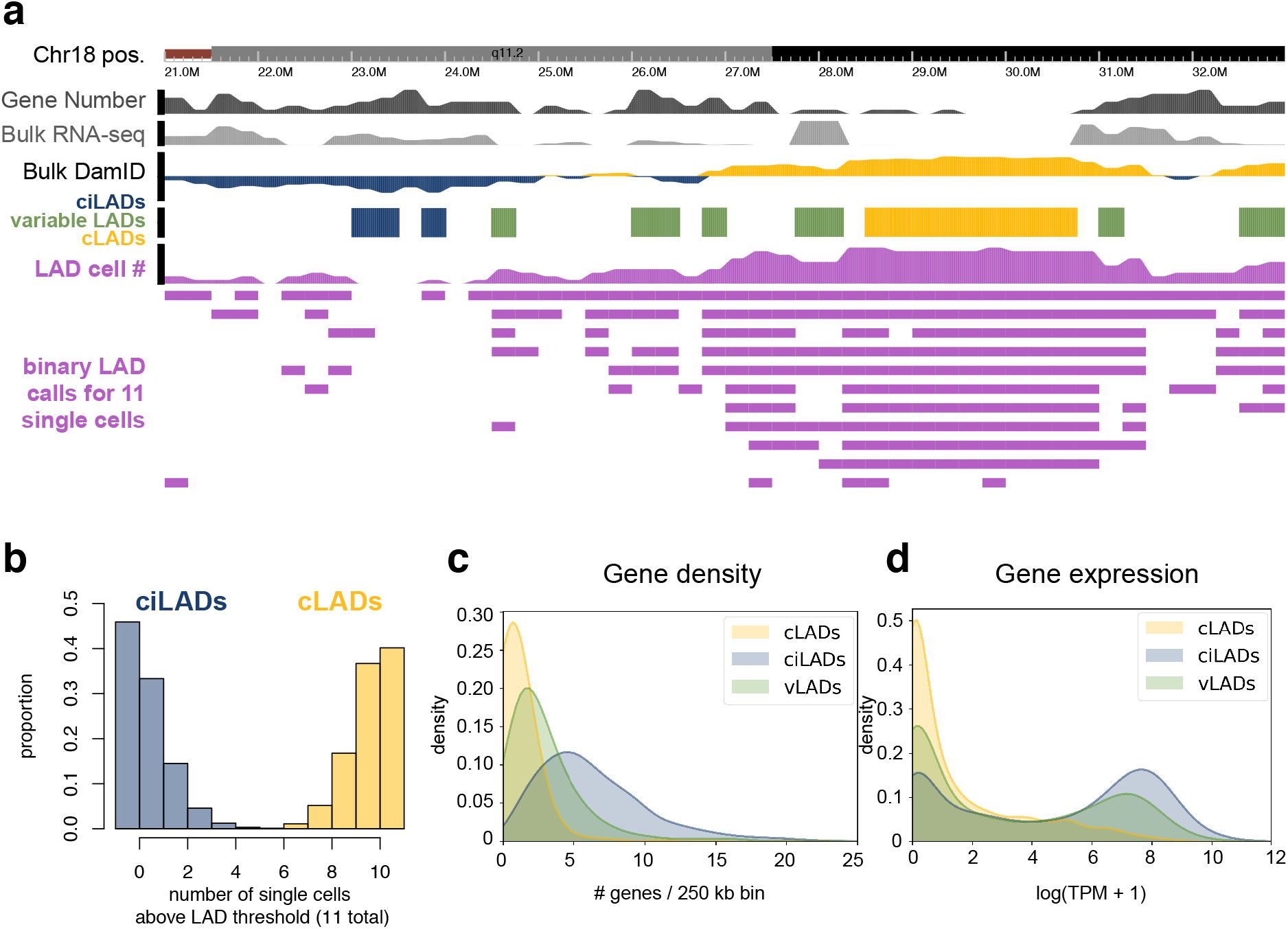
Defining variable LADs in HEK293T cells. (**a**) A browser screenshot from Chr18:21-33 Mb. The first track shows the chromosome ideogram and coordinates. The second track reports the number of Refseq genes falling in each bin. The third track reports the mean Transcripts Per Million (TPM) value for each gene within each bin from bulk RNA-seq data from untreated HEK293T cells. The fourth track reports the log_2_FoldChange values from bulk Dam-LMNB1:Dam-only sequencing data. The fifth track indicates the positions of the control cLAD (gold) and ciLAD (blue) sets as well as the positions of regions called as variable LADs using the single-cell sequencing data generated here (green). The sixth track shows the number of single cells (out of 11) in which each bin is called as an LAD. Below that, the positions of all bins called as LADs are indicated, with one row per cell. (**b**) Distribution of the number of single cells (out of 11) in which each bin is called as an LAD for all 250 kb bins genome-wide, separately for each of the control sets of cLADs or ciLADs. (**c-d**) Distributions of the number of genes (**c**) or mean TPM per gene (**d**) per 250 kb bin for each of the sets of cLADs, ciLADs, or variable LADs.

To classify bins confidently as variable LADs, we aimed to rule out the possibility that sampling error could explain the observed intermediate number of LAD-classified cells in these regions, given the range of error rates within individual cells. Among bins called as LADs in 4-7 cells, we computed the joint probability of observing that number of cells under two null models: one consisting of true positives and false negatives, and one consisting of true negatives and false positives (see Methods). We selected only the subset of bins with low p-values (p<10^−8^) under both null models, providing high confidence that these variable LAD regions are truly variable between cells (Figure 3a). We hypothesized that these stringently defined regions, which comprise 13% of the genome, would be more gene rich and have higher gene expression than cLADs, given their dynamic positioning in cells. Indeed, these variable LADs show intermediate gene density and bulk gene expression levels compared to the control sets of cLADs and ciLADs (Figure 3c-d), consistent with these regions being variably active within different cells.

### μDamID enables cell-cell comparisons based on imaging and sequencing data

μDamID enables the joint analysis of the nuclear localization and sequence identity of protein-DNA interactions within each cell and between cells. Because the nuclear localization of LADs is well characterized, one could generate and test hypotheses about the sequencing data given the imaging data for each cell in this study. For example, cells expressing Dam-only show fluorescence throughout the center of the nucleus, and indeed their coverage profiles show little difference in coverage between known cLADs and ciLADs (Figure 4). Moreover, Dam-LMNB1 cells with visible rings and low fluorescence in the nuclear interior tend to show well-separated cLAD and ciLAD coverage distributions (Figure 4). One anomalous Dam-LMNB1 cell (cell #7) was selected for having bright fluorescence throughout the nucleus, and its sequencing data confirm that it appears to have increased coverage in ciLADs, appearing like an intermediate between the Dam-only and Dam-LMNB1 coverage signatures (Figure 4). Dam-LMNB1 is likely overexpressed in that cell, causing it to accumulate high background levels of methylation throughout the nucleus.

**Figure 4.**
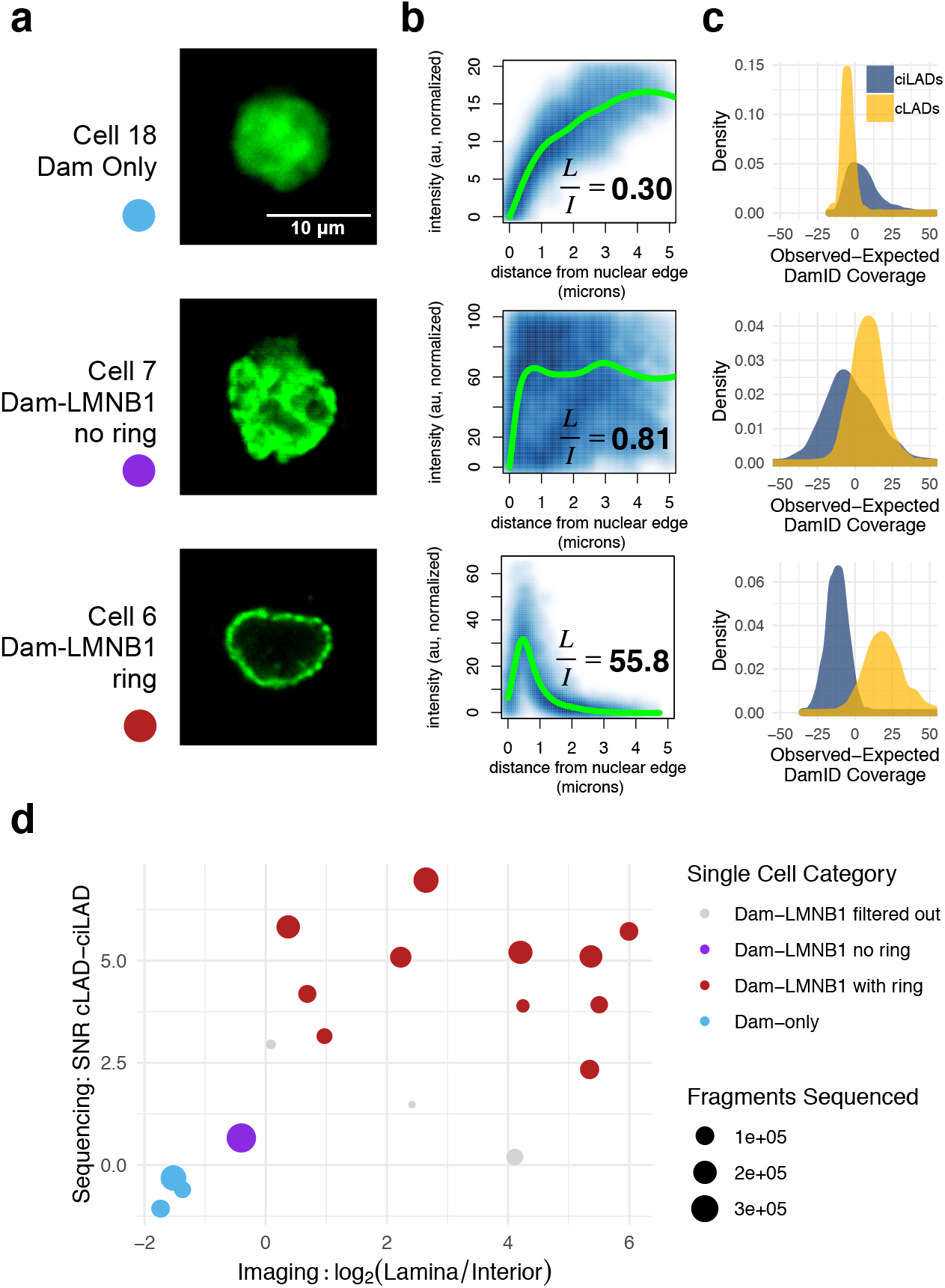
Joint imaging and sequencing analysis with μDamID. (**a**) Confocal fluorescence microscopy images of ^m6^A-Tracer GFP signal from 3 cells: one expressing Dam-only, one expressing Dam-LMNB1 but showing high interior fluorescence, and one expressing Dam-LMNB1 and showing the expected ring-like fluorescence at the nuclear lamina. (**b**) Normalized pixel intensity values plotted as a function of their distance from the nuclear edge (blue), with a fitted loess curve overlaid (green). Ratios of the mean normalized pixel intensities in the Lamina (<1 micron from the edge) versus the Interior (>3.5 microns from the edge) are printed on each plot. (**c**) DamID sequencing coverage distributions for each of the cLAD or ciLAD control sets (as in Figure 2c). (**d**) scatterplot showing sequencing versus imaging metrics for each cell, with point size indicating the number of unique DpnI fragments sequenced for that cell. The x axis reports the log_2_ ratio of the Lamina:Interior mean intensity ratio for each cell. The y axis reports the log_2_ of the Signal-to-Noise Ratio (SNR) computed from the sequencing data for each cell (effectively the difference in means between cLADs and ciLADs divided by the standard deviation of ciLAD coverage).

To quantify these observations across all cells, for each image we generated an averaged GFP intensity profile plot as a function of the distance from the edge of the nuclear lamina (Figure 4b). Using these profiles, we computed the ratio of mean GFP intensity at the nuclear lamina compared to the nuclear interior, which is small for the Dam-only cells and cell #7, and large for the Dam-LMNB1 cells. Then, we compared these imaging ratios to a computed sequencing signal-to-noise ratio (SNR) for each cell, a measure of how well separated the cLAD and ciLAD coverage distributions are (see Methods and Figure 4d). The Dam-only and Dam-LMNB1 cells can be readily separated on either axis, with cell #7 appearing intermediate on both axes. Overall, these data add additional confidence that the sequenced areas correspond to the fluorescing areas of the nucleus, providing two useful measures of chromatin organization within single cells.

## Conclusions

We have demonstrated the use of an integrated microfluidic device for single-cell isolation, imaging, and sorting, followed by DamID. This system enables the acquisition of paired imaging and sequencing measurements of protein-DNA interactions within single cells, giving a readout of both the ‘geography’ and identity of these interactions in the nucleus. Specifically, we tested the device by mapping well-characterized interactions between DNA and proteins found at the nuclear lamina, providing a measure of genome regulation and 3D chromatin organization within the cell, and recapitulating similar maps in other cell types. This technology could be applied to study many other types of protein-DNA interactions in single cells, and it could be combined with other sequencing and/or imaging modalities to gather even richer information from each cell. For example, the nuclear localization of specific proteins such as heterochromatin-associated proteins or nucleolus-associated proteins can be visualized by fluorescent tagging, then DamID can be used to sequence and identify nearby genomic regions. Recent advances allow for simultaneous DamID and transcriptome sequencing in single cells (Rooijers et al. 2019), and this device could be adapted for similar multi-omic protocols as well. Further improvements to the DamID protocol may also improve its sensitivity and specificity. Finally, one can increase the number of cells processed per batch by scaling up the design and incorporating features like multiplexed valve control and automated image processing and sorting.

## Materials and Methods

### Cell transfection and harvesting

HEK293T cells (CRL-3216, ATCC, Manassas, VA; validated by microsatellite typing, at passage number <10) were seeded in 24-well plates at 50000 cells per well in 0.5 ml media (DMEM plus 10% FBS). The next day, cells were transfected using FuGene HD transfection reagent according to their standard protocol for HEK293 cells (Promega, Madison, WI). DNA plasmids were cloned in Dam-negative *E. coli* to reduce sequencing reads originating from plasmid. Dam-LMNB1 and ^m6^A-Tracer plasmids were obtained from Bas van Steensel (from Kind et al. 2013); Dam-LMNB1 was modified to replace GFP with mCherry and to produce a Dam-only version; their sequences are available as supplementary information (see link to GitHub below in Data Availability section). 250 ng Dam construct DNA plus 250 ng ^m6^A-Tracer DNA were used per well. As controls to validate transfection, additional wells were left untransfected, transfected with ^m6^A-Tracer only, or transfected with Dam construct only. The following day, successful transfection was validated by widefield fluorescence microscopy, seeing GFP signal in wells containing ^m6^A-Tracer, and mCherry signal in all wells containing Dam construct only. Cells were harvested 72 hours after transfection. 20 hours before harvesting, the media was replaced and 0.5 μl Shield-1 ligand (0.5 mM stock, Takara Bio USA, Inc., Mountain View, CA) was added to each well to stabilize protein expression. Cells transfected with Dam-LMNB1 were inspected by fluorescence microscopy to look for the characteristic signal at the nuclear lamina, indicating proper expression and protein activity. To harvest the cells and prepare them for loading on the device, the cells were washed with PBS, then incubated at room temperature with 1X TrypLE Select (ThermoFisher Scientific, Waltham, MA) for 5 minutes to dissociate them from the plate. Cells were pipetted up and down to break up clumps, then centrifuged at 300xg for 5 minutes, resuspended in PBS, centrifuged again, and resuspended in 500 μl Pick Buffer (50mM Tris-HCl pH 8.3, 75mM KCl, 3mM MgCl2, 137mM NaCl), achieving a final cell concentration of roughly 500,000 cells per ml. Cells were passed through a 40 μm cell strainer before loading onto the device.

### Confocal imaging

Fluorescence confocal imaging of cells was performed in the trapping region using an inverted scanning confocal microscope with a 488 nm Ar/Kr laser (Leica, Germany) for excitation, with a bandpass filter capturing backscattered light from 500-540 nm at the primary photomultiplier tube (PMT), with the pinhole set to 1 Airy unit, with a transmission PMT capturing widefield unfiltered forward-scattered light, and with a 63X 0.7 NA long-working-distance air objective with a correction collar, zoomed by scanning 4X. Gain and offset values were set automatically for one cell and identical microscope settings were used to image all cells. The focal plane was positioned in the middle of each nucleus, capturing the largest-circumference cross-section, and final images were averaged over 10 frames to remove noise. The 3 cells expressing Dam-only that were sequenced in this study were imaged with a widefield CCD camera. Other Dam-only cells were imaged with confocal microscopy and showed similar relatively homogenous fluorescence throughout the nucleus, and never the distinct ‘ring’ shape found in Dam-LMNB1 expressing cells (Kind et al. 2013; Supplementary Figure 5). No image enhancement methods were used prior to quantitative image processing. Images in Figures 1 and 4 have been linearly thresholded to diminish background signal.

### Mold fabrication

Molds for casting each layer were fabricated on silicon wafers by standard photolithography. Photomasks for each layer were designed in AutoCAD and printed at 25400 DPI (CAD/Art Services, Inc., Bandon, Oregon). The mask for the thick layer, in this case the flow layer to make push-up valves, was scaled up in size uniformly by 1.5% to account for thick layer shrinkage. A darkfield mask was used for features made out of negative photoresist: the filters on the flow layer and the entire control layer; a brightfield mask was used for all flow layer channels, which were made out of positive photoresist (mask designs available on GitHub; see Data Availability section below). 10 cm diameter, 500 μm thick test-grade silicon wafers (item #452, University Wafer, Boston, MA) were cleaned by washing with 100% acetone, then 100% isopropanol, then DI water, followed by drying with an air gun, and heating at 200°C for 5 minutes.

To make the control layer mold, SU-8 2025 negative photoresist (MicroChem Corp., Westborough, MA) was spin-coated to achieve 25 μm thickness (7 s at 500 rpm with 100 rpm/s ramp, then 30 s at 3500 rpm with 300 rpm/s ramp). All baking temperatures, baking times, exposure dosages, and development times followed the MicroChem data sheet. All baking steps were performed on pre-heated ceramic hotplates. After soft-baking, the wafer was exposed beneath the darkfield control layer mask using a UV aligner (OAI, San Jose, CA). After post-exposure baking and development, the mold was hard-baked at 150°C for 5 minutes.

To make the flow layer mold, first the filters were patterned with SU-8 2025, which was required to make fine, high-aspect-ratio filter features that would not re-flow at high temperatures. SU-8 2025 was spin-coated to achieve 40 μm thickness (as above but with 2000 rpm final speed) and processed according to the MicroChem datasheet as above, followed by an identical hard-bake step. Next, AZ 40XT-11D positive photoresist (Integrated Micro Materials, Argyle, TX) was spin-coated on top of the SU-8 features to achieve 20 μm thickness across the wafer (as above but with 3000 rpm final speed). All baking temperatures, baking times, exposure dosages, and development times followed the AZ 40XT-11D data sheet. After development, the channels were rounded by reflowing the photoresist, achieved by placing the wafer at 65°C for 1 min, then 95°C for 1 min, then 140°C for 1 min followed by cooling at room temperature. In our experience, reflowing for too long, or attempting to hard-bake the AZ 40XT-11D resulted in undesirable ‘beading’ of the resist, especially at channel junctions. Because it was not hard-baked, no organic solvents were used to clean the resulting mold. Any undeveloped AZ 40XT-11D trapped in the filter regions was carefully removed using 100% acetone applied locally with a cotton swab.

### Soft lithography

Devices were fabricated by multilayer soft lithography (Unger et al. 2000). On-ratio 10:1 base:crosslinker RTV615A PDMS (Momentive Performance Materials, Inc., Waterford, NY) was used for both layers, and layer bonding was performed by partial curing, followed by alignment, then full curing (Lai et al. 2019). To prevent PDMS adhesion to the molds, the molds were silanized by exposure to trichloromethlysilane (Sigma-Aldrich, St. Louis, MO) vapor under vacuum for 20 min. PDMS base and crosslinker were thoroughly mixed by an overhead mixer for 2 minutes, then degassed under vacuum for 90 minutes. Degassed PDMS was spin-coated on the control layer mold (for the ‘thin layer’) to achieve a thickness of 55 μm (7 s at 500 rpm with 100 rpm/s ramp, then 60 s at 2000 rpm with 500 rpm/s ramp), then placed in a covered glass petri dish and baked for 10 minutes at 70°C in a forced-air convection oven (Heratherm OMH60, Thermo Fisher Scientific, Waltham, MA) to achieve partial curing. The flow layer mold (for the ‘thick layer’) was placed in a covered glass petri dish lined with foil, and degassed PDMS was poured onto it to a depth of 5 mm. Any bubbles were removed by air gun or additional degassing under vacuum for 5 minutes, then the thick layer was baked for 19 minutes at 70°C. Holes were punched using a precision punch with a 0.69 mm punch tip (Accu-Punch MP10 with CR0420275N19R1 punch, Syneo, Angleton, TX). The thick layer was peeled off the mold, cut to the edges of the device, and aligned manually under a stereoscope on top of the thin layer, which was still on its mold. The layers were then fully cured and bonded together by baking at 70°C for 120 min. After cooling, the devices were peeled off the mold, and the inlets on the thin layer were punched. The final devices were bonded to 1 mm thick glass slides, which were first cleaned by the same method as used for silicon wafers above, using oxygen plasma reactive ion etching (20 W for 23 s at 285 Pa pressure; Plasma Equipment Technical Services, Brentwood, CA), followed by heating at 100°C on a ceramic hot plate for 5 minutes.

### Device and control hardware setup

Devices were pneumatically controlled by a solenoid valve manifold (Pneumadyne, Plymouth, MN). Each three-way, normally open solenoid valve switched between a regulated and filtered pressure source inlet at 25 psi (172 kPa) or ambient pressure to close or open microfluidic valves, respectively. Solenoid valves were controlled by the KATARA control board and software (White and Streets 2018). Most operational steps were carried out on inverted microscopes using 4-10X objectives. For incubation steps, the device was placed on a custom-built liquid-cooled thermoelectric temperature control module (TC-36-25-RS232 PID controller with a 36 V 16 A power source and two serially connected VT-199-1.4-0.8P TE modules and an MP-3022 thermistor; TE technologies, Traverse City, MI) controlled by a new KATARA software module (to be made available on github). A layer of mineral oil was applied between the chip and the temperature controller to improve thermal conductivity and uniformity. A stereoscope was used to monitor the chip while on the temperature controller.

To set up each new device, each pneumatic valve was connected to one control inlet on the microfluidic device by serially connecting polyurethane tubing (3/32” ID, 5/32” OD; Pneumadyne) to Tygon tubing (0.5 mm ID, 1.5 mm OD) to >4 cm PEEK tubing (0.25 mm ID, 0.8 mm OD; IDEX Corporation, Lake Forest, IL). Solenoid valves were energized to de-pressurize the tubing and the tubing was primed by injecting water using a syringe connected to the end of the PEEK tubing, then the primed PEEK tubing was inserted directly into each punched inlet hole on the device. Solenoid valves were de-energized to pressurize the tubing until all control channels on the device were fully dead-end filled, then each microfluidic valve was tested and inspected by switching on and off its corresponding solenoid valve. All valves were opened and the device was passivated by filling all flow-layer channels with syringe-filtered 0.2% (w/w) Pluronic F-127 solution (P2443; MilliporeSigma, St. Louis, MO) from the reagent inlet and incubating at room temperature for 1 hour. The device was then washed by flowing through 0.5 ml of ultra-filtered water, followed by air to dry it.

### Device operation

Initially, all chamber valves and reagent inlet valves were closed. Gel-loading pipette tips were used to inject reagents and cells into the flow channels. To prepare the device for operation, pick buffer was injected into the reagent inlet and pressurized at 5-10 psi to dead-end fill the reagent inlet channels. Negative controls were generated by injecting pure pick buffer into one of the holding chambers before trapping and sorting cells into the other lanes. 50 μl of cell suspension was then loaded into a gel-loading pipette tip, and injected directly into the cell inlet. A high-precision pressure regulator was used to load the single-cell suspension at 1 psi (7 kPa). Cells were observed in the filter region with brightfield and epifluorescence using a 10X objective to identify candidate cells. These were then tracked through the device until they approached the trapping chamber for an empty lane. To trap a candidate cell, the device’s peristaltic pump was operated at 1 Hz to deliver that cell to the trap region. The trap valves (above and below the trap region; see Figure S1) were closed and the cell was imaged with scanning confocal microscopy as described above. If the cell was rejected after imaging, the trap valves were opened and it was flushed to the waste outlet. Otherwise, the cell was injected into the holding chamber by dead-end filling. This process was repeated to fill each lane with single cells for DamID. To test background DNA levels, we filled the final lane with only cell suspension buffer. Nearly undetectable levels of amplified DNA were recovered from these lanes.

After filling all 10 lanes, the reagent inlet and cell trapping channels were flushed with 0.5 ml of water, which exited through the waste outlet and the cell inlet, to remove any remaining Pick buffer or cell debris, then air dried. The same washing and drying was repeated between each reaction step. To inject reagents for each reaction of the DamID protocol, the trap valves were closed, the reagent channels were dead-end filled with freshly prepared and syringe-filtered reagent, then the reagent inlet valves and the valves for the necessary reaction chambers were opened, and each lane was dead-end filled individually to prevent any possible cross-contamination. Reaction contents are described in Table 1.

**Table 1.** Reaction buffers and conditions

| Reaction Stage | Buffer | Incubation |
| --- | --- | --- |
| Trapping & Holding | Pick Buffer:<br>50mM Tris-HCl pH 8.3<br>75mM KCl, 3mM MgCl<sub>2</sub><br>137mM NaCl | RT |
| Lysis | 10mM TRIS acetate pH 7.5<br>10mM magnesium acetate 50mM potassium acetate<br>0.67% Tween-20<br>0.67% Igepal<br>0.67 mg/ml proteinase K | 42 °C for 4 hours<br>then 80 °C for 10 min |
| Digestion | mix 7μl 10X Cutsmart buffer<br>1 μl DpnI (New England Biolabs, Ipswich, MA)<br>62 μl H<sub>2</sub>O | 37 °C for 4 hours<br>then 80 °C for 20 min |
| Ligation | mix 6 μl 10X NEB T4 ligase buffer<br>1 μl DamID adapter stock at 25 μM<br>0.2 μl NEB T4 ligase at 400 U/μl<br>21.8 μl H<sub>2</sub>O<br>1 μl 2% w/v Tween-20 | 16 °C overnight<br>then 65 °C for 10 min |
| PCR | from Takara Clontech Advantage 2 kit:<br>mix 5 μl 10X PCR buffer<br>1 μl dNTPs at 10 mM each<br>1 μl polymerase mix<br>0.63 μl DamID primer<br>21.37 μl H<sub>2</sub>O<br>1 μl 2% Tween-20 | See Table 2 |

After filling all lanes, reagents were mixed by actuating the chamber valves at 5 Hz for 5 minutes. At the PCR step, rotary mixing was achieved by using the chamber valves to make a peristaltic pump driving fluid around the full reaction ring. For each reaction step, the device was placed on the thermal controller and reactions were with times and temperatures described in Table 1. PCR thermocycling conditions are described in Table 2. To ensure adequate hydration during PCR, all valves were pressurized. Amplified DNA was flushed out of each lane individually using purified water from the reagent inlet, collected into a gel loading tip placed in the lane outlet to a final volume of 5 μl then transferred to a 0.2 ml PCR strip tube.

**Table 2.** PCR thermocycling conditions

| PCR Step | Incubation |
| --- | --- |
| 1 | 68 °C for 10 min |
| 2 | 94 °C for 1 min |
| 3 | 65 °C for 5 min |
| 4 | 68 °C for 15 min |
| 5 | 94 °C for 1 min |
| 6 | 65 °C for 1 min |
| 7 | 68 °C for 10 min |
| 8 | Go to step 5 (x 3) |
| 9 | 94 °C for 1 min |
| 10 | 65 °C for 1 min |
| 11 | 68 °C for 2 min |
| 12 | Go to step 9 (x 22) |
| 13 | Hold 10 °C |

### Oligonucleotides

>AdRt
CTAATACGACTCACTATAGGGCAGCGTGGTCGCGGCCGAGGA
>AdRb
TCCTCGGCCG
>AdR_PCR
NNNNGTGGTCGCGGCCGAGGATC

To anneal DamID adapter (from Vogel et al. 2007): mix equal volumes of 50 μM AdRt and 50 μM AdRb in a microcentrifuge tube, then fully submerge it in a beaker of boiling water, and allow the water to equilibrate to room temperature slowly.

### Quality control, library preparation, and sequencing

Samples were diluted to 10 μl total volume and two replicates of qPCR were performed using the DamID PCR primer to estimate DNA quantities relative to the pick-buffer-only negative control (1 μl DNA per replicate in 10 μl reaction volume). We also used 1 μl of sample to measure DNA concentration using a Qubit fluorometer with a High-Sensitivity DNA reagent kit (quantitative range 0.2 ng – 100 ng; ThermoFisher Scientific). Samples with the lowest Ct values and highest Qubit DNA measurements were selected for library preparation and sequencing. Library preparation was carried out using an NEBNext Ultra II DNA Library Prep Kit for Illumina (NEB E7645) with dual-indexed multiplex i5/i7 oligo adapters. Size selection was not performed; PCR was carried out for 9 cycles. Libraries were quantified again by Qubit and size profiled on a TapeStation 4200 with a D5000 HS kit (Agilent, Santa Clara, CA), then mixed to achieve equimolar amounts of each library. DNA was sequenced on an Illumina MiniSeq with a 150-cycle high output kit, to produce paired 75 bp reads, according to manufacturer instructions (Illumina, San Diego, CA). Roughly 13 million read pairs were obtained.

### Bulk DamID

Genomic DNA was isolated from ~3.7 x 10^6^ transfected HEK293T cells using the DNeasy Blood & Tissue kit (Qiagen) following the protocol for cultured animal cells with the addition of RNase A. The extracted gDNA was then precipitated by adding 2 volumes of 100% ethanol and 0.1 volume of 3 M sodium acetate (pH 5.5) and storing at −20 °C for 30 minutes. Next, centrifugation for 30 minutes at 4 °C, >16,000 x g was performed to spin down the gDNA. The supernatant was removed, and the pellet was washed by adding 1 volume of 70% ethanol. Centrifugation for 5 minutes at 4 °C, >16,000 x g was performed, the supernatant was removed, and the gDNA pellets were air-dried. The gDNA was dissolved in 10 mM Tris-HCl pH 7.5, 0.1 mM EDTA to 1 μg/μl, incubating at 55 °C for 30 minutes to facilitate dissolving. The concentration was measured using Nanodrop.

The following DpnI digestion, adaptor ligation, and DpnII digestion steps were all performed in the same tube. Overnight DpnI digestion at 37 °C was performed with 2.5 μg gDNA, 10 U DpnI (NEB), 1X CutSmart (NEB), and water to 10 μl total reaction volume. DpnI was then inactivated at 80 °C for 20 minutes. Adaptors were ligated by combining the 10 μl of DpnI-digested gDNA, 1X ligation buffer (NEB), 2 μM adaptor dsAdR, 5 U T4 ligase (NEB), and water for a total reaction volume of 20 μl. Ligation was performed for 2 hours at 16 °C and then T4 ligase was inactivated for 10 minutes at 65 °C. DpnII digestion was performed by combining the 20 μl of ligated DNA, 10 U DpnII (NEB), 1X DpnII buffer (NEB), and water for a total reaction volume of 50 μl. The DpnII digestion was 1 hour at 37 °C followed by 20 minutes at 65 °C to inactivate DpnII.

Next, 10 μl of the DpnII-digested gDNA was amplified using the Clontech Advantage 2 PCR Kit with 1X SA PCR buffer, 1.25 μM Primer Adr-PCR, dNTP mix (0.2 mM each), 1X PCR advantage enzyme mix, and water for a total reaction volume of 50 μl. PCR was performed with an initial extension at 68 °C for 10 minutes; one cycle of 94 °C for 1 minute, 65 °C for 5 minutes, 68 °C for 15 minutes; 4 cycles of 94 °C for 1 minute, 65 °C for 1 minute, 68 °C for 10 minutes; 21 cycles of 94 °C for 1 minute, 65 °C for 1 minute, 68 °C for 2 minutes. Post-amplification DpnII digestion was performed by combining 40 μl of the PCR product with 20 U DpnII, 1X DpnII buffer, and water to a total volume of 100 μl. The DpnII digestion was performed for 2 hours at 37 °C followed by inactivation at 65 °C for 20 minutes. The digested product was purified using QIAquick PCR purification kit.

The purified PCR product (1 μg brought up to 50 μl in TE) was sheared to a target size of 200 bp using the Bioruptor Pico with 13 cycles with 30”/30” on/off cycle time. DNA library preparation of the sheared DNA was performed using NEBNext Ultra II DNA Library Prep Kit for Illumina.

### Bulk DamID, comparing Dam mutants

Bulk DamID for comparing the wild-type allele and V133A mutant allele was performed as outlined in the Bulk DamID methods section with the following modifications. Genomic DNA was extracted from ~ 2.4 x 10^5^ transfected HEK293T cells. A cleanup before methylation-specific amplification was included to remove unligated Dam adapter before PCR. The Monarch PCR & DNA Cleanup Kit with 20 μl DpnII-digested gDNA input and an elution volume of 10 μl was used. Shearing with the Bioruptor Pico was performed for 20 total cycles with 30”/30” on/off cycle time. Paired-end 2 x 75 bp sequencing was performed on an Illumina NextSeq with a mid output kit. Approximately 3.8 million read pairs per sample were obtained.

### Bulk RNA-seq

RNA was extracted from ~1.9 x 10^6^ transfected HEK293T cells using the RNeasy Mini Kit from Qiagen with the QIAshredder for homogenization. RNA library preparation was performed using the NEBNext Ultra II RNA Library Prep Kit for Illumina with the NEBNext Poly(A) mRNA Magnetic Isolation Module. Paired-end 2 x 150 bp sequencing for both DamID-seq and RNA-seq libraries was performed on 1 lane of a NovaSeq S4 run. Approximately 252 million read pairs were obtained for each DamID-seq sample, and roughly 64 million read pairs for each RNA sample. Adapters were trimmed using trimmomatic (v0.39; Bolger et al. 2014; ILLUMINACLIP:adapters-PE.fa:2:30:10 LEADING:3 TRAILING:3 SLIDINGWINDOW:4:15 MINLEN:36, where adapters-PE.fa is:

>PrefixPE/1
TACACTCTTTCCCTACACGACGCTCTTCCGATCT
>PrefixPE/2
GTGACTGGAGTTCAGACGTGTGCTCTTCCGATCT).

Transcript quantification was performed using Salmon (Patro et al. 2017) with the GRCh38 transcript reference. Differential expression analysis was performed using the voom function in limma (Ritchie et al. 2015). Differential expression was called based on logFC significantly greater than 1 and adjusted p-value < 0.01.

### DamID sequence processing and analysis

Bulk and single-cell DamID reads were demultiplexed using Illumina’s BaseSpace platform to obtain fastq files for each sample. DamID and Illumina adapter sequences were trimmed off using trimmomatic (v0.39; Bolger et al. 2014; ILLUMINACLIP:adapters-PE.fa:2:30:10 LEADING:3 TRAILING:3 SLIDINGWINDOW:4:15 MINLEN:20, where adapters-PE.fa is:

>PrefixPE/1
TACACTCTTTCCCTACACGACGCTCTTCCGATCT
>PrefixPE/2
GTGACTGGAGTTCAGACGTGTGCTCTTCCGATCT).
>Dam
GGTCGCGGCCGAGGA
>Dam_rc
TCCTCGGCCGCGACC

). Trimmed reads were aligned to a custom reference (hg38 reference sequence plus the Dam-LMNB1 and ^m6^A-Tracer plasmid sequences) using BWA-MEM (v0.7.15-r1140, Li 2013).

Alignments with mapping quality 0 were discarded using samtools (v1.9, Li et al. 2009). The hg38 reference sequence was split into simulated DpnI digestion fragments by reporting all intervals between GATC sites (excluding the GATC sites themselves), yielding 7180359 possible DpnI fragments across the 24 chromosome assemblies. The number of reads overlapping each fragment was counted using bedtools (v2.28; Quinlan et al. 2010). For single-cell data, the number of DpnI fragments with non-zero coverage was reported within each non-overlapping bin in the genome (11512 total 250 kb bins). For bulk data, the number of read pairs overlapping each 250 kb bin was reported. The same exact pipeline was applied to the raw reads from Kind et al. 2015 in aggregate. RefSeq gene positions were downloaded from the UCSC Genome Browser and counted in each bin. For bulk data, Dam-LMNB1 vs DamOnly enrichment was computed using DEseq2 in each 250 kb bin (Love et al. 2014). For single-cell data, the expected background coverage in each bin was computed as *n*(*m*/*t*), where *n* is the number of unique fragments sequenced from that cell, *m* is the number of bulk Dam-only read pairs mapping to that bin, and *t* is the total number of mapped bulk Dam-only read pairs. Single-cell normalization was computed either as a ratio of observed to expected coverage (for browser visualization and comparison to bulk data), or as their difference (for classification and coverage distribution plotting). Positive and negative control sets of cLAD and ciLAD bins were defined as those with a bulk Dam-LMNB1:Dam-only DEseq2 p-value smaller than 0.01/11512, that intersected published cLADs and ciLADs in other cell lines (Lenain et al. 2017), and that were among the top 1200 most differentially enriched bins in either direction (positive or negative log fold change for cLADs and ciLADs, respectively). Integer normalized coverage thresholds for LAD/iLAD classification were computed for each cell to maximize accuracy on the cLAD and ciLAD control sets. Signal-to-noise ratios were computed for each cell using the normalized coverage distributions in the cLAD and ciLAD control sets as (*μ*_cLAD_ − *μ*_ciLAD_)/*σ*_ciLAD_. Statistical analyses and plots were made in R (v3.5.2) using the ggplot2 (v3.1.0), gplots (v3.0.1.1), and colorRamps (v2.3) packages. Browser figures were generated using the WashU Epigenome Browser (Li et al. 2019).

### Image processing

Images were processed in R (v3.5.2) and plots were produced using the reshape2 (v1.4.3), SDMTools (v1.1-221.1), spatstat (v1.59-0), magick (v2.0), ggplot2 (v3.1.0), and ggbeeswarm (v0.6) packages. Grayscale images were converted to numeric matrices and edge detection was performed using Canny edge detection using the image_canny function in magick, varying the geometry parameters manually for each cell. The center of mass of all edge points was obtained, and all edge points were plotted in Cartesian coordinated with this center of mass as the origin. Noise was removed by removing points with a nearest neighbor more than 2 microns away. Edge point coordinates were converted to polar coordinates, and the farthest points from the origin in each 10 degree arc were reported. Within each 10 degree arc, all pixel intensities from the original image within the edges of the nucleus were reported as a function of their distance from the farthest edge point in that arc to make Figure 4b. For each cell a loess curve (span 0.3) was fitted to the data after subtracting the minimum intensity value within 3.5 microns of the edge. The Lamina:Interior ratio was computed as the ratio of mean intensity of pixels within 1 micron of the edge to the mean intensity of pixels more than 3.5 microns from the edge, after subtracting the minimum value of the loess curve for that cell.

## Author contributions

NA and AMS conceived of and designed the study and the microfluidic device. NA and AL fabricated and optimized operation of the device. AM performed bulk cell experiments and data processing, and NA performed all other experiments, analysis, and pneumatic/thermoelectric hardware construction. JAW developed the microfluidic control platform and thermal cycling software, with minor modifications by NA. NA wrote the manuscript with contribution from AM and AMS. AMS supervised the study.

## Acknowledgements

We would like to thank Anushka Gupta, Gabriel Dorlhiac, Zoë Steier, Adam Gayoso, Tyler Chen, Xinyi Zhang, and Carolina Rioz-Martinez for their helpful feedback on this work. We are grateful to Carolyn de Graaf for providing us with LAD coordinates, to Bo Huang for providing guidance and materials, and to Bas van Steensel for providing us with plasmids. Nicolas Altemose is supported by a Howard Hughes Medical Institute Gilliam Fellowship for Advanced Study. This work was supported by the National Institute of General Medical Sciences of the National Institutes of Health [Grant Number R35GM124916]. Aaron M. Streets is a Chan Zuckerberg Biohub Investigator.

## Conflicts of interest

The authors declare no competing interests.

## Data availability

Sequencing data are available on FigShare: https://doi.org/10.6084/m9.figshare.8856368.

Imaging data are available on FigShare: https://doi.org/10.6084/m9.figshare.8940245.

Analysis code, control software, device design files, and plasmid sequences are available on

GitHub: https://github.com/altemose/microDamID.

## Supplementary Figures

**Supplementary Figure 1.**
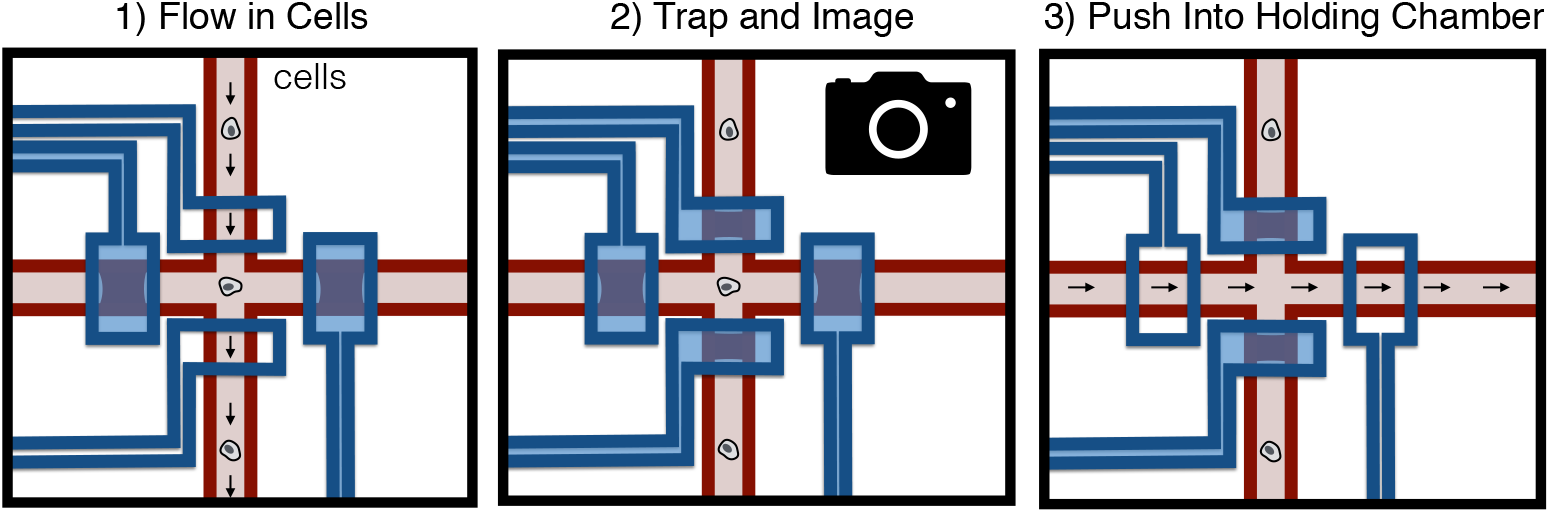
Illustration of cell trapping procedure. Cells are driven through the device by peristaltic pumping or pressure-driven flow. Valves are actuated to confine the cell in the trapping region, where it is imaged, and if selected, is pushed by dead-end filling into a holding chamber to the right of the trapping region.

**Supplementary Figure 2.**
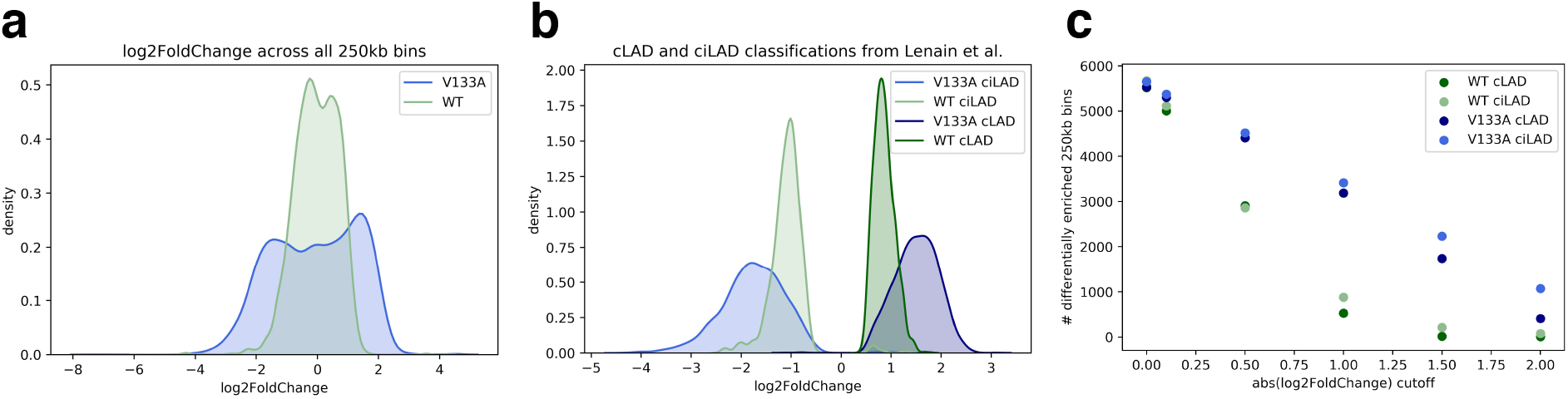
Comparison of wild-type and V133A Dam alleles. (**a**) Kernel density estimate of log_2_FoldChange from DESeq2 differential enrichment analysis of Dam-LMNB1 coverage compared to Dam-only as reference. With V133A, more extreme log_2_FoldChange values are observed with greater separation between the positive and negative log_2_FoldChange peaks. In other words, compared to wild-type, the V133A Dam-LMNB1 and Dam-only signals are more distinct. (**b**) Kernel density estimate of log_2_ Fold Change, with cLAD/ciLAD classification from Lenain et al. 2017 indicated, shows greater separation for cLAD and ciLAD signal with V133A. (**c**) V133A has higher sensitivity than WT, with more differentially enriched regions at each log_2_FoldChange threshold for calling significant differential enrichment.

**Supplementary Figure 3.**
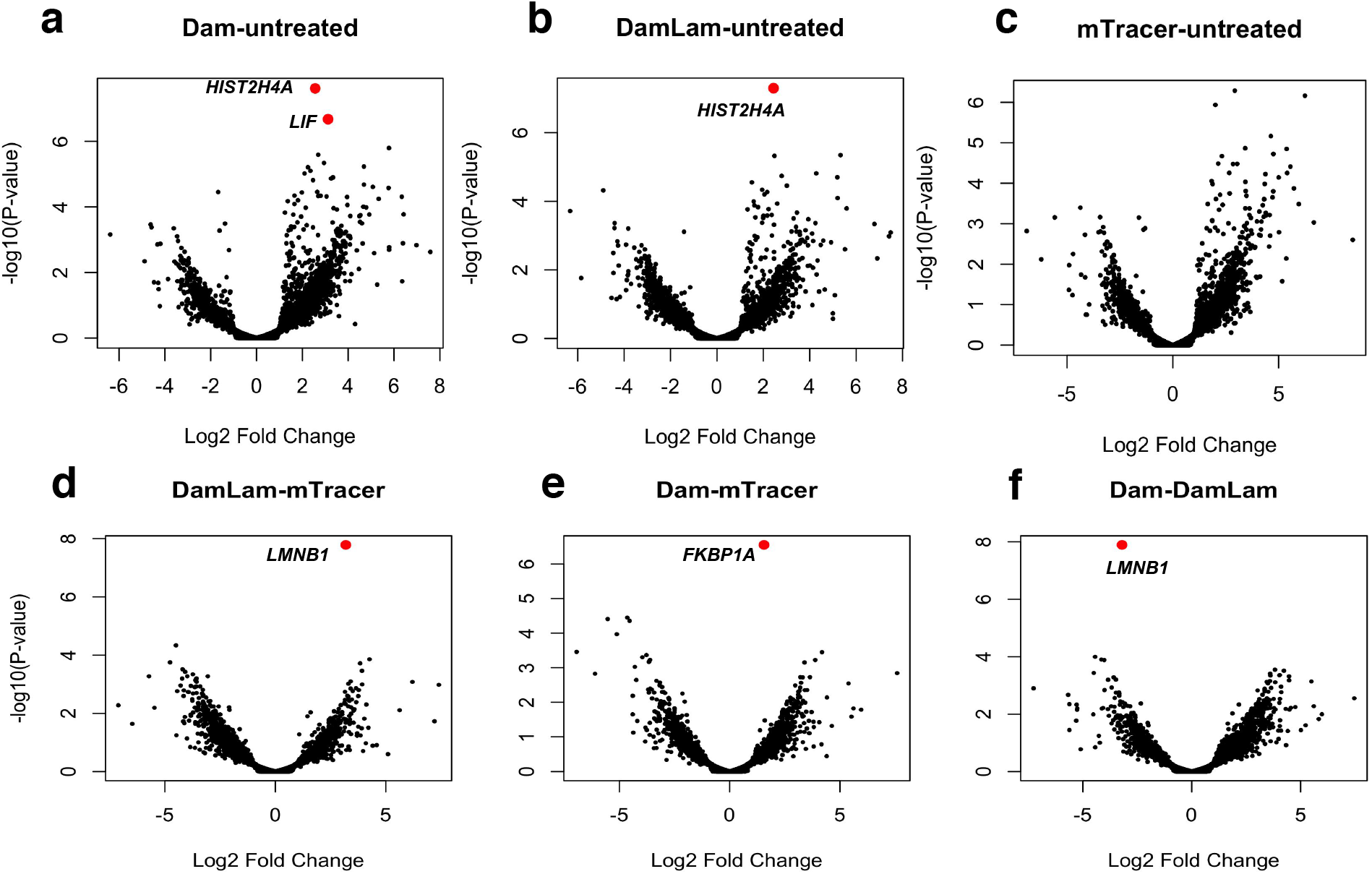
Volcano plots from differential gene expression analysis for RNA from bulk HEK293T cells transfected with Dam, *Dam-LMNB1*, ^m6^A-Tracer, or no treatment control. Significantly differentially expressed genes (logFC significantly > 1 and adjusted p-value < 0.01) are indicated in red. Differentially expressed genes compared to no treatment control are *HIST2H4A* and *LIF* for Dam, *HIST2H4A* for Dam-LMNB1, and no genes for ^m6^A-Tracer. When comparing Dam to ^m6^A-Tracer, the only differentially expressed gene is *FKBP1A*, which is expected given the mutated FKBP1A-derived destabilization domain tethered to Dam in our construct. When comparing Dam-LMNB1 to ^m6^A-Tracer, the only differentially expressed gene is *LMNB1*, which is again expected given *LMNB1* is expressed from the *Dam-LMNB1* construct itself.

**Supplementary Figure 4.**
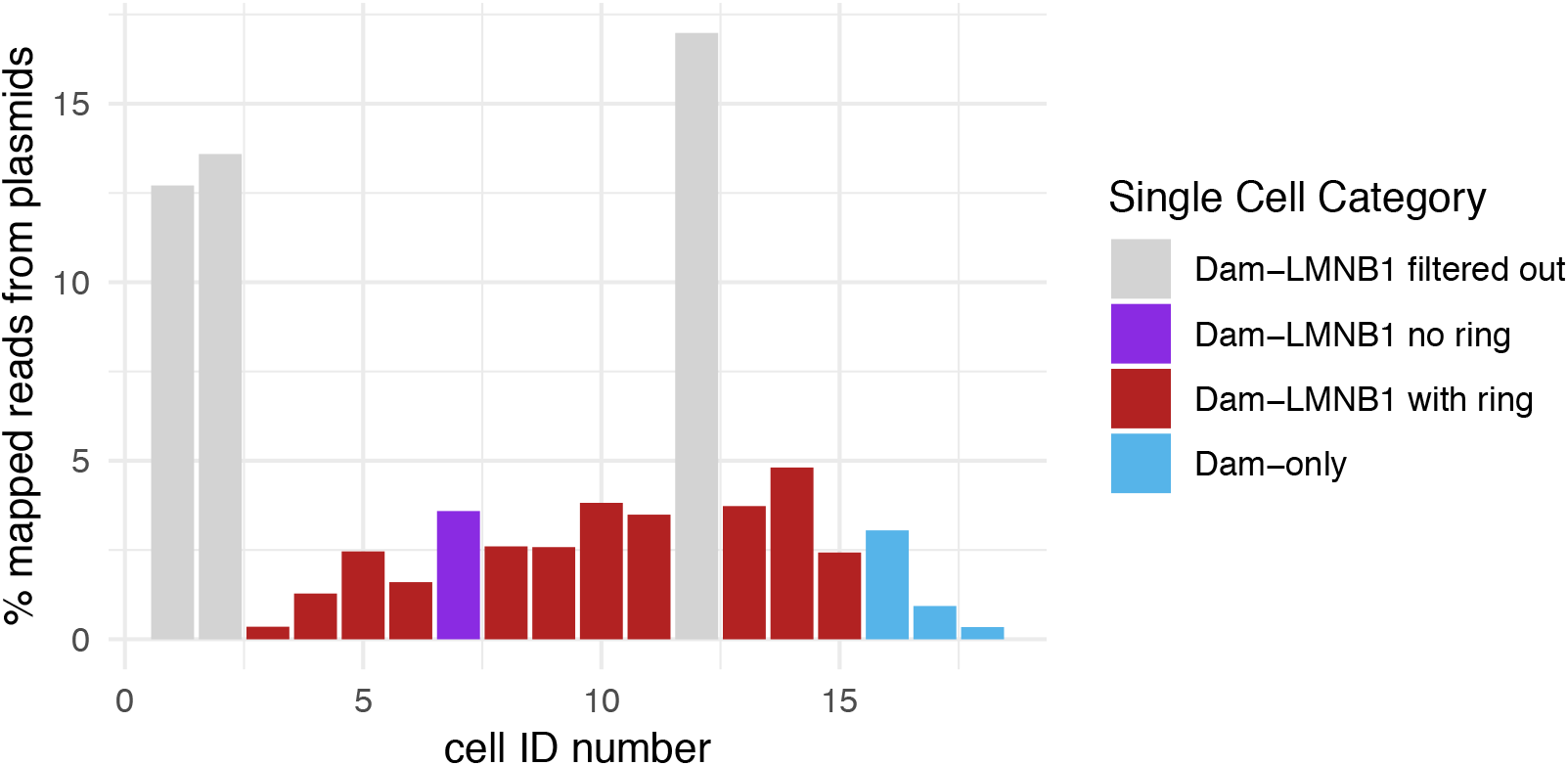
Percentage of mapped sequencing reads mapping to plasmid sequences for each single cell. Cells 1, 2, and 12 were filtered out due to their high plasmid DNA content.

**Supplementary Figure 5.**
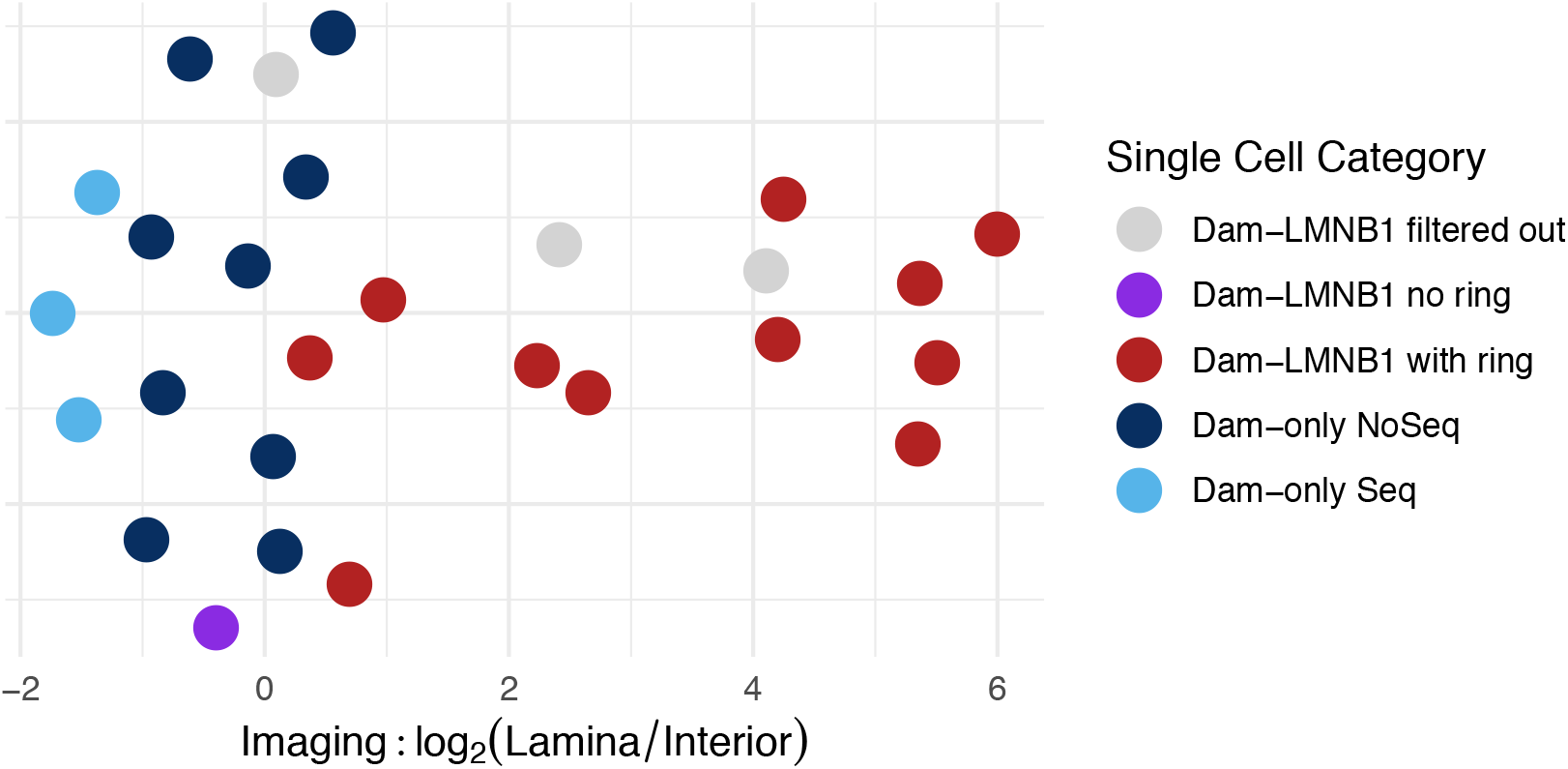
Lamina:Interior mean pixel intensity ratios for all cells. Imaging ratios are reported for each cell as in Figure 4d. Dark blue points represent Dam-only cells that were imaged by confocal microscopy but not sequenced, compared to the light blue points representing the Dam-only cells that were imaged by widefield microscopy and sequenced.

